# Enhancing Light-Sheet Fluorescence Microscopy Illumination Beams through Deep Design Optimization

**DOI:** 10.1101/2023.11.29.569329

**Authors:** Chen Li, Mani Ratnam Rai, Yuheng Cai, H. Troy Ghashghaei, Alon Greenbaum

## Abstract

Light sheet fluorescence microscopy (LSFM) provides the benefit of optical sectioning coupled with rapid acquisition times for imaging of tissue-cleared specimen. This allows for high-resolution 3D imaging of large tissue volumes. Inherently to LSFM, the quality of the imaging heavily relies on the characteristics of the illumination beam, with the notion that the illumination beam only illuminates a thin section that is being imaged. Therefore, substantial efforts are dedicated to identifying slender, non-diffracting beam profiles that can yield uniform and high-contrast images. An ongoing debate concerns the employment of the most optimal illumination beam; Gaussian, Bessel, Airy patterns and/or others. Comparisons among different beam profiles is challenging as their optimization objective is often different. Given that our large imaging datasets (∼0.5TB images per sample) is already analyzed using deep learning models, we envisioned a different approach to this problem by hypothesizing that we can tailor the illumination beam to boost the deep learning models performance. We achieve this by integrating the physical LSFM illumination model after passing through a variable phase mask into the training of a cell detection network. Here we report that the joint optimization continuously updates the phase mask, improving the image quality for better cell detection. Our method’s efficacy is demonstrated through both simulations and experiments, revealing substantial enhancements in imaging quality compared to traditional Gaussian light sheet. We offer valuable insights for designing microscopy systems through a computational approach that exhibits significant potential for advancing optics design that relies on deep learning models for analysis of imaging datasets.

## 1. Introduction

Current advancements in imaging can be attributed to two parallel research efforts: First, the improvement of optical system design and components to achieve superior properties, such as imaging speed and depth, signal-to-noise ratio, and more [1–7]; Second, the utilization of advanced post-processing algorithms to enhance image quality and potentially streamline the data extraction phase [8–14]. The latter approach has gained popularity with the development of deep learning techniques due to the fact that it does not require alternations to existing optical setups, and its ability to faithfully mimic human analysis of the data [15–19]. More recently, there is an ongoing effort to merge these two research streams and jointly optimize both the imaging system and the deep learning network [20–25]. This approach involves incorporating the physical model into the mathematical framework of deep learning, which we refer to as “Deep Design” (DD).

Here, we will demonstrate the potential of DD in the context of Light-Sheet Fluorescence Microscopy (LSFM), which necessitates the incorporation of a very complex optical model into the optimization process. LSFM has become an indispensable imaging tool in the life science community due to its high acquisition speed and optical sectioning capabilities [26–31]. In LSFM, the illumination path is separated and orthogonal to the detection path. A thin plane of light is used to illuminate the tissue, and fluorophores within the excitation volume are excited and their emitted light is collected using a wide-field detection system which is oriented perpendicularly to the light-sheet illumination axis [32,33]. LSFM image quality depends on various factors. Lateral resolution is influenced by fluorescence wavelength and detection objective numerical aperture (NA), while axial resolution and signal-to-noise ratio are broadly influenced by the illumination pattern shape and detection objective NA [31,34]. As a result, the image quality of LSFM is not uniform across the Field of View (FOV), often resulting in less-than-ideal datasets. Figure 1A illustrates the problem: a classical light sheet beam is tightly focused on the center, resulting in clear and sharp images. However, due to diffraction, the edges of specimen are illuminated by a broader beam, leading to noisy images with reduced axial resolution.

**Figure 1.**
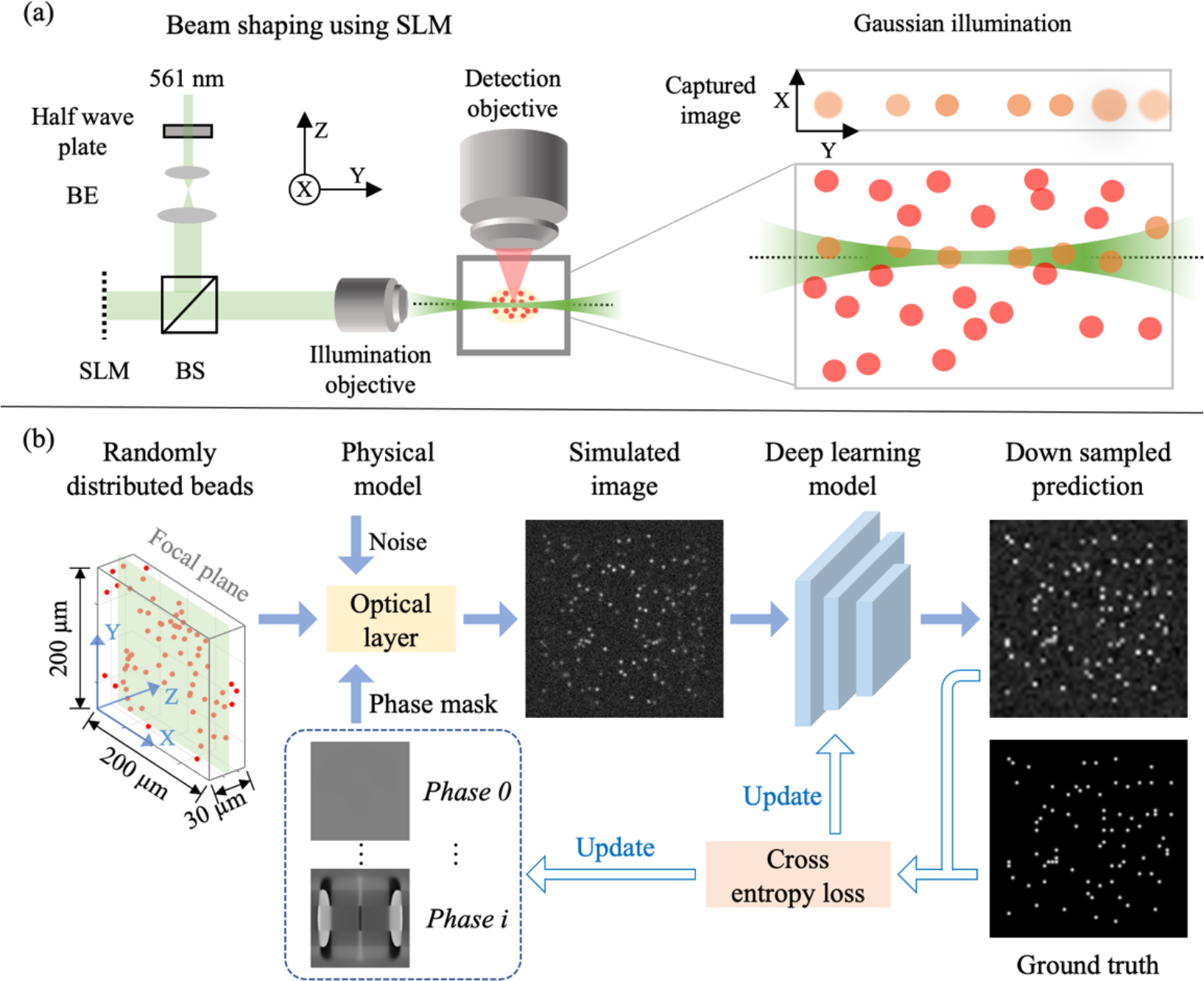
Deep Design (DD) approach to improve optical sectioning and image quality in LSFM. **(a)** A simplified optical setup that contains a spatial light modulator (SLM) to control the properties of the illumination beam. The close-up highlights a challenge encountered when employing a Gaussian beam in light sheet fluorescence microscopy (LSFM): diffraction causes the illumination beam to widen at the edges. Consequently, out-of-focus beads are unintentionally illuminated, introducing noise into the image. BS – beam splitter; BE – beam expander. **(b)** Joint optimization scheme. The locations of randomly distributed beads will be fed into a physical optical layer to generate simulated images. The prediction network outputs a 2D downsized image to predict the position of beads in the focal plane. The deep learning network and input phase mask are simultaneously updated based on the loss function.

To mitigate the issue of non-uniform image quality in LSFM, there is a continuous search for methods to generate non-diffracting and pseudo-nondiffracting beams that maintain a consistent illumination profile across the FOV while confining the illumination profile to a narrow plane. Past studies have explored different illumination patterns such as Bessel [35,36] and Airy beams [37,38], which then combined with LSFM to improve image contrast and resolution. However, debates continue about the ideal illumination beam since quantitative comparisons are challenging due to differences in input pupil structures and NA requirements [34,39].

Here we showcase an alternative conceptual perspective on the problem by engineering the LSFM illumination beam using DD. DD operates within <u>two core layers</u> (Fig. 1b): (1) <u>A Physical Modeling Layer</u> that functions as a simulator, generating a 2D image that a LSFM would capture when confronted with a random distribution of fluorescence emitters within a defined volume and a specific phase mask. The phase mask delineates the illumination pattern within the volume. (2) <u>A Deep Learning Layer</u> that contains a deep learning model which acquires knowledge of the illumination pattern’s shape and detects objects. From simulations, our DD generated a novel LSFM illumination beam that enhanced optical sectioning – which we hereafter refer to as the butterfly beam (the phase mask resembles a butterfly). This innovative illumination beam is the outcome of optimizing millions of variables within the phase mask. Subsequently, we conducted <u>experimental validation</u>, revealing that the “butterfly beam” exhibits significantly narrower edges within the FOV compared to a Gaussian beam with an identical spatial light modulator (SLM) aperture size. This result underscores the significance of integrating the physical model into the deep design framework to advance the performance of optical system design. It constitutes a paradigm shift in the field of optical system design when data is automatically analyzed by deep learning models.

## 2. Methods

Our premise was that enhanced optical sectioning in LSFM will lead to better object detection. This assumption dictated our optimization target in simulation, i.e., enhancing the detection of fluorescent emitters within the focal plane while effectively suppressing signals originating from out-of-focus regions. The architecture of our end-to-end network was structured around two key components. First, a differentiable optical simulation layer was incorporated, featuring a trainable phase-only mask positioned placed directly against the illumination objective (Fig. 1a). This component can change the shape of the illumination beam, affording a unique opportunity to optimize the shape of the light-sheet for enhanced image quality. The second pivotal component was a convolutional neural network (CNN), renowned for its prowess in feature extraction and pattern recognition. Within our framework, CNN played an indispensable role by predicting the precise locations of in-focus signals within the simulated images. In this section we provide the physical and mathematical framework of the problem.

### 2.1 Optical simulation layer

The optical simulation layer functioned as follows: 1) Initially, we created a random distribution of densely packed beads (3 μ*m* in diameter) within a 3D volume measuring 200×200×30 μ*m*^3^. 2) We then simulated the light intensity that interacted with each individual bead within the volume. This 3D intensity distribution undergone changes according to the phase mask pattern. It is important to note that due to the non-ideal nature of the generated light sheet, beads located outside the focal plane of the detection objective were also illuminated. Subsequently, we generated the recorded image by considering the noise produced by out-of-focus beads.

#### 2.1.1 Simulating the intensity of the variable illumination beam across the 3D volume

To simulate the illumination conditions [40], we first modeled the intensity emitted from the laser as a gaussian beam defined as:

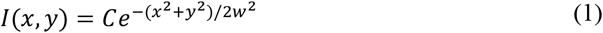

Where *x*, and *y* represented spatial coordinates, *w* was the waist of the input beam, and *C* was an arbitrary constant. To simplify our simulation, we assumed that the field of the input beam was multiplied by a defined phase pattern ℬ (*x, y*) and placed directly against the imaging lens. The result was then focused on the sample. The complex field at the focal plane of the illumination lens can be defined as:

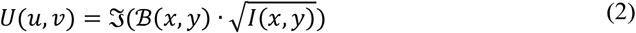

Where ℑ was the Fourier transform, *u, and v* were the spatial coordinates in the focal plane and *F* was the illumination lens focal distance. *u and v* were defined as *u* = *f*_*x*_λ*F* and *v* = *f*_*y*_λ*F*, where λ was the illumination wavelength, and *f*_*x*_, *and f*_*y*_ were the spatial frequencies of the Fourier transform. Next, to propagate the field along the volume of excitation, for an arbitrary distance L, we used the angular spectrum method of propagation. The complex field was defined as

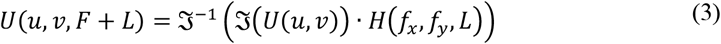

Where *H* was the transfer function of propagation through free space.

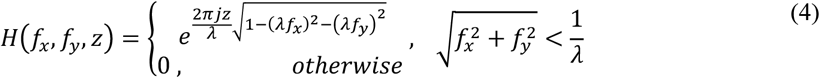

After generating the spatial profile of the beam, the beam was computationally dithered up and down to form the light-sheet. The entire process was modeled in our DD approach, and ℬ (*x, y*) was the variable phase mask which determined the optical property of the generated beam. ℬ(*x, y*) was jointly optimized with the detection network.

#### 2.1.2. Synthetic image formation

Once the intensity that impinged on each bead in the volume was known, its effect on the detected image needed to be determined. To this end, we calculated the bead point spread function (PSF) as a function of its distance from the detection objective focal plane (z distance) [41]. The defocused incoherent PSF was defined as follows:

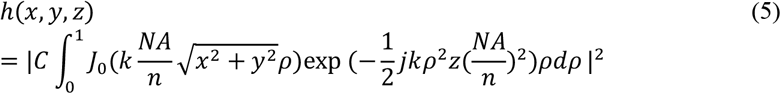

Where *J*_*0*_ was the Bessel function of the *1*^st^ kind, *n* = 1 was the refractive index, *C* = 1 was an arbitrary normalization constant, *NA* = 0.6 was the numerical aperture, λ = 561 *nm* was wavelength, *k* = 2π/λ was the wave number.

To create a complete synthetic image, we aggregated the individual bead images produced by variable PSFs and combined them with the light sheet intensity. The result was a synthesized image that provided a comprehensive representation of the observed scene. To ensure the fidelity and realism of our synthesized image, we introduced Poisson noise into the image data. This noise accounted for various factors, including the offset of the image sensor and random background noise. This step was crucial for simulating real-world conditions and improving the accuracy of our results. In Figure 1b, a physical model was presented to illustrate the process of generating a synthetic image.

### 2.2 Joint-optimization of the phase mask and object detection network

We used a rudimentary convolutional neural network to detect the location of the beads, which contained only 6 layers of Resnet with short-cut connections. State-of-the-art object detection networks did not fit into our limited GPU memory (32 GB) given the incorporation of the complex optical simulation layer. The end-to-end deep learning pipeline was illustrated in Figure 1b. The physical model took the random beads locations (*x, y, z*), and given a phase mask it generated synthetic images (pixel size 1 × 1 μm^2^). The network component consisted of the repeated convolution layers with 3×3 kernel size, followed by leaky-Relu activation function and batch normalization. Max-pooling was applied to perform down samplings to reduce computational cost. The output of the network was a 2D grid with pixel size of 4 × 4 μm^2^ which indicted the position of in-focus beads. The detailed network structure is shown in Figure S1.

Because the optical model was differentiable, the network parameters and phase masks can be jointly optimized simultaneously. The joint-network was implemented in Python 3.7 with PyTorch-1.12.0 Deep Learning Framework and trained on one single Nvidia Tesla V100-32GB GPU on the UNC Longleaf cluster for ∼72 hours. The prediction of in-focus beads was treated as a classification problem, and the binary cross entropy with logits loss function was used. The loss function was calculated between the ground truth of in-focus 2D grid of beads and the output grid of the network. The learning rate was set to 1e-2 with Adam optimizer. The pixel size of phase mask and synthetic image was 1 × 1 μm^2^, and the size of phase mask and synthetic image was set to 500 × 500 μm^2^ and 200 × 200 μm^2^, respectively. Simulated beads were randomly distributed in the 200 × 200 × 20 μm^3^ space. Details about the parameter settings can be found in the source code.

### 2.3 Custom-built light sheet design

LSFM was inspired by previous designs [42–45], and the basic setup and complete list of components can be found in our recent studies [29,43]. Here we duplicated the setup and added a SLM to reshape the illumination beam. Briefly, a Gaussian beam emitted from a continuous-wave laser (Coherent; OBIS LS 561-50) served as our illumination source. The beam passed through a half wave plate that was mounted on rotating fixture. We expanded the beam and used a scanning galvo system (Cambridge Technology; 6215H) to introduce dithering up and down of the illumination beam, which consequently generated the light sheet. This scanning galvo system was conjugated onto a spatial light modulator (Medowlark E19X12 series) to optimize the phase within the beam path. The beam, now modulated by the SLM, underwent focusing through a lens (100 mm), with a slit positioned at the front focal plane of the lens. The purpose of the slit was to eliminate the 0*th* order (unmodulated light) while collecting only the modulated light. Further demagnification was achieved through a combination of a 2X lens (Keyence BZ-PF10P 0.1 NA) and a 10X lens (Olympus RMS10X-PF 0.30), resulting in a slenderer beam. Subsequently, this refined beam was projected onto a cuvette (Thorlabs CV10Q35FAE) filled with DBE, serving as the chamber for imaging tissue-cleared samples. The interface between the lens and cuvette was filled with immersion oil to minimize aberrations. Additionally, to correct for any illumination aberrations, the Gaussian beam aberrations were recorded and corrected using the SLM, and the corrections were used as constant biases on the SLM pattern. The placement of the sample within the cuvette was facilitated by a customized ASI-3D stage, allowing for precise 3D adjustments during imaging. To capture the fluorescent signal, a 10X objective lens (10X Mitutoyo Plan Apo 0.28 NA) was employed, and an emission filter (AVRO; FF01-593/40-25) was inserted into the detection path to eliminate unwanted signals. The detected signal was then collected by a tube lens (ASI; TL180-MMC), followed by a CMOS camera (Hamamatsu, C13440-20CU). A graphical user interface was written in MATLAB (2019b), which was used during image acquisition.

### 2.4 Sample preparation

Two mouse brains [46] and four prairie vole brains were subjected to pretreatment involving methanol, immunolabeling, and clearing [47]. Postnatal day 30 mouse brain samples were stained using chicken anti-GFP (with Alexa Fluor 647 as the secondary antibody) and rabbit anti-RFP (with Cy3 as the secondary antibody). Whereas, Postnatal day 60 vole brain were stained using mouse anti-oxytocin (with Alexa Fluor 647 as the secondary antibody) and rabbit anti-vasopressin (with Alexa Fluor 555 as the secondary antibody). The harvesting of all animals was conducted in accordance with the regulations and approval of the Institutional Animal Care and Use Committee (IACUC) at North Carolina State University.

## 3. Results

### 3.1 Simulation results – The butterfly beam improves object detection

We trained our end-to-end deep learning model from random distributed beads in 3D space, starting with an initial phase mask of all zeros, which generated a conventional Gaussian light-sheet. Through each iteration, the phase mask was updated together with the object detection network to improve detection accuracy. From simulation results, the DD method produced a phase mask that resulted in a light-sheet with less diffraction along the propagation axis (Fig. 2a), after 50 epochs. Figure 2b illustrates the calculated full width half maximum of the butterfly (optimized) and gaussian beams. The full width at half maximum (FWHM) was determined by collapsing the beam along the dithering direction. A Gaussian fit was then applied to the averaged beam, and the FWHM was calculated based on the standard deviation (SD) as FWHM = 2.355×SD. The butterfly beam shows a much narrower width at the edges of the FOV, while slightly sacrificing the middle of the FOV. A detailed cross-section of the Gaussian and our butterfly beam in simulation is provided in Figure S2. Simulation of the butterfly beam produced cleaner images at the edges (i.e., less background noise) in comparison with the Gaussian beam (Fig. 2a, blue boxes).

**Figure 2.**
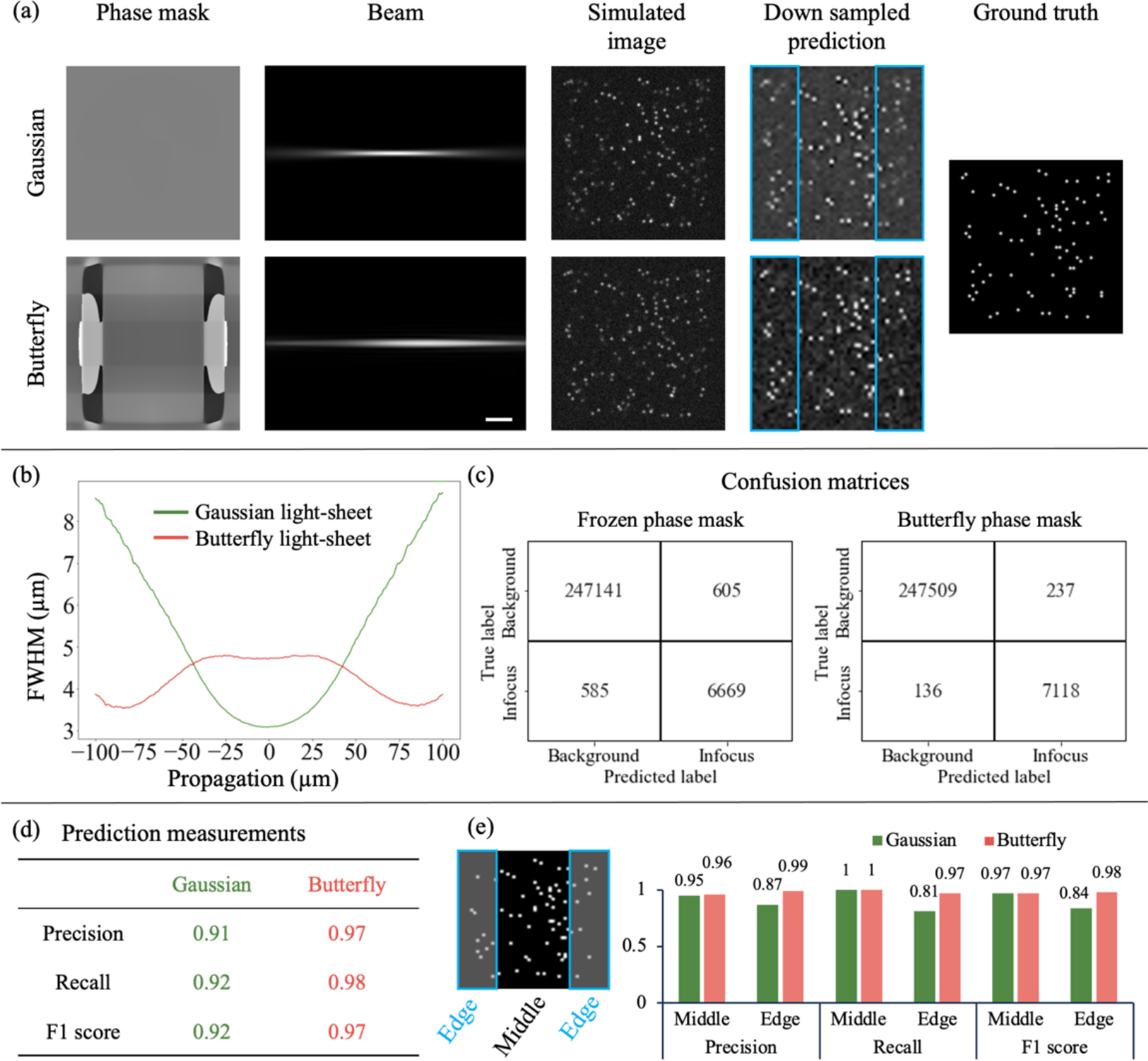
Simulation of DD-mediated optimization. (a) Optimized phase mask to improve bead detection accuracy. Compared with a Gaussian beam (flat phase mask), the butterfly beam exhibits better image contrast on the edges (see inside the blue rectangles area), rendering a better prediction. **(b) Beam profile comparison**. Full Width at Half Maximum (FWHM) for Gaussian and butterfly beams plotted against the direction of propagation. The graph illustrates that the butterfly beam exhibits reduced FWHM at the edges, while the Gaussian beam is narrower at the center of the Field of View. **(c-d) Network performance**. A comparison of classification metrics (confusion matrix and prediction measurement) between frozen and butterfly phase scenarios. The optimized phase mask demonstrates improved bead detection capability and achieves higher scores across the evaluated metrics. **(e) Comparison between middle and edge area**. The entire FOV is divided into middle and edge areas. The metrics are calculated separately, and the butterfly beam provides better performance on the areas corresponding to the edge.

The output of the DD approach was also an object detection network, which is tailored to the butterfly beam illumination profile. Therefore, we validated that the improvement in image quality at the edges of the FOV was translated to improvement in the downstream analysis (Figs. 2c-e). For an unbiased comparison, our baseline\gold standard network was trained from scratch on images that were produced solely by a Gaussian beam. We refer to this network as the frozen phase mask network since it was not updated from a uniform phase during the training. Figure. 2c shows the confusion matrix for two classes: background and in-focus beads. The overall F1-score (indicating class-wise performance) of the butterfly and Gaussian beam was 0.97, and 0.92 respectively (Fig. 2d). Since the butterfly beam was narrower at the edges of the FOV, we also compared the object detection results between the center and the edges of the FOV (Fig. 2e). We expected to see that the object detection in conjunction with the butterfly beam will outperform the Gaussian beam on the edges, which was confirmed by simulation. For instance, at the edge of the field-of-view, the F1-score of the butterfly and Gaussian beam was 0.98, and 0.84 respectively. To evaluate DD robustness, we have run DD under different noise levels and initial guess conditions - initial guess refers to the initialization value of the phase mask when initiating DD optimization. Remarkably, in all tested cases the butterfly beam pattern has emerged as the most optimal illumination beam (Fig. S3-S4). As an additional quality check for our DD approach, we also tested it under perturbation experiment (Fig. S5). For instance, when we focused the beam waist away from the center of the FOV, the network repositioned it by producing a lens-like pattern (Fig. S5). Qualitatively, the same butterfly beam shape was observed under these perturbation experiments.

### 3.2 Experimental results – the butterfly beam shape and profile are similar to the simulation

Following, we conducted experimental tests on the properties of the butterfly beam. Figure 3a reveals that, consistent with the simulation, the experimentally measured FWHM of the butterfly beam was wider at the center, measuring 11.93 μm, compared to both edges of the FOV, for instance, 9.35 μm (at 400 μm) or 10.93 μm (at -600 μm). This contrasted sharply with the Gaussian beam (featuring a constant phase mask on the SLM), where the FWHM was narrower at the center, measuring 8.33 μm, compared to both FOV edges, such as 14.67 μm (at 400 μm) or 22.24 μm (at -600 μm). Conceptually, these numerical values closely paralleled our expectations from simulation (Fig. 2).

**Figure 3.**
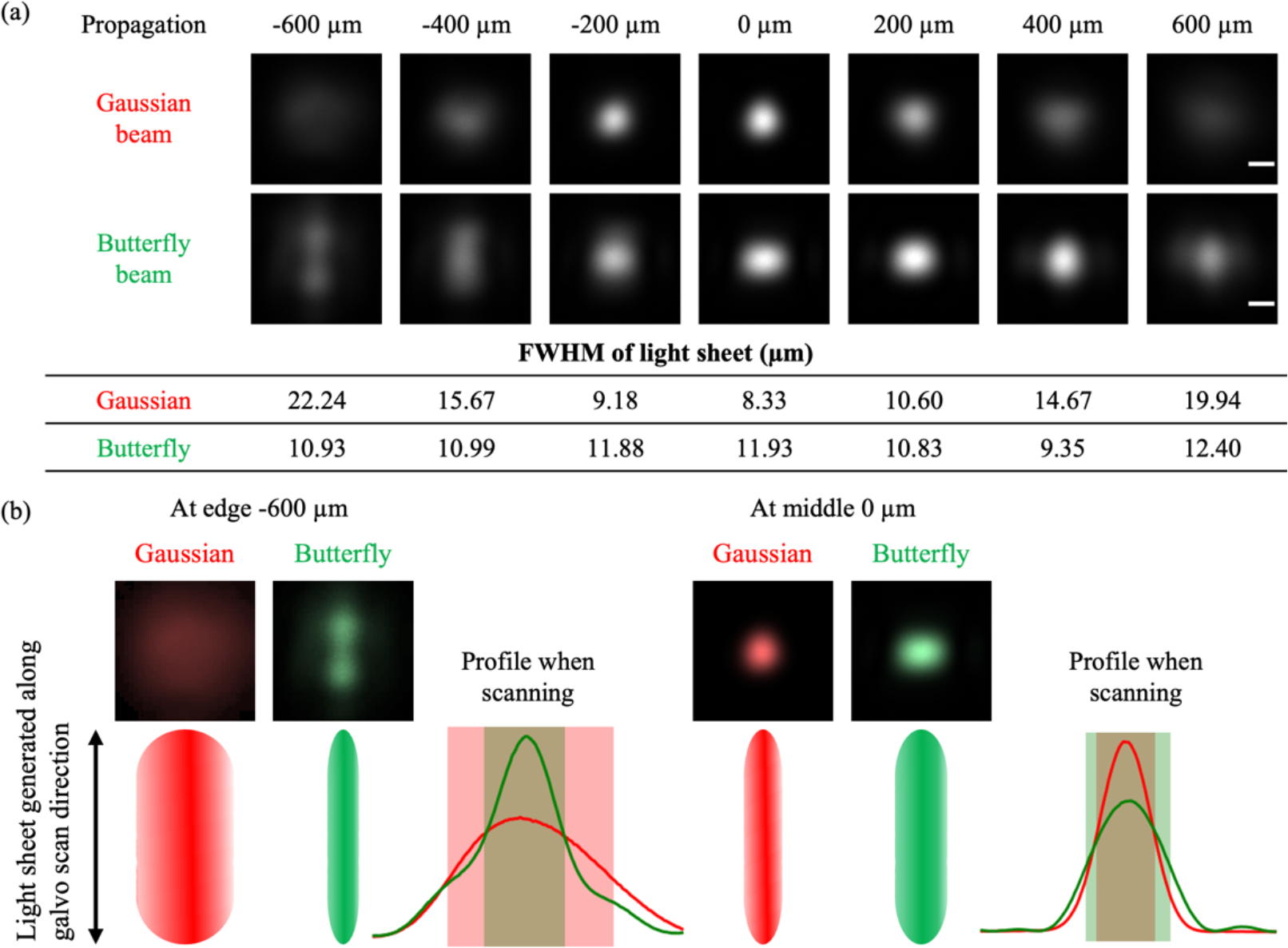
Experimental results - characterization of the butterfly beam. **(a)**. In the top panels, images depict the static profile of Gaussian and butterfly beams along the beam propagation. The propagation distance of 0 μm corresponds to the focal point of the excitation objective, where the Gaussian beam reaches its minimal waist. As the light sheet is generated by dithering the static beam up and down, light sheet thickness was measured by summing the values along the columns. Notably, experimental measurements of the beam were conducted using a single lens after the spatial light modulator, outside the immersion chamber. The scale bar is 10 μm. **(b)** Schematic depicting the formation of the light sheet through the up-and-down dithering of the static beam. The line profile along the scanning direction compares the Gaussian beam (in red) with the optimized/butterfly beam (in green). To maintain a narrow profile at the edges of the field of view, the deep design optimization elongated the profile in the direction of the scan.

Please note that the experimental setup underwent modifications from the simulation. This adjustment was necessitated by the need to utilize very short distances during the simulation to replicate high numerical aperture conditions within the constraints of limited GPU memory. Within these brief distances (e.g., 2 mm), the integration of optical components in an experimental setup proved unfeasible. Consequently, the primary disparities between the simulation and experimental setup encompassed: 1) The physical size of the SLM exceeded the dimensions of the phase mask in the simulation. 2) For the generation of the butterfly beam, the SLM was positioned in the back focal plane of a 1’’ lens with a 125 mm focal distance (Thorlabs; AC254-125-A-ML). 3) The SLM featured constant phase patterns for aberration compensation and a linear phase mask to reject the 0st order.

Figure 3b illustrates the utilization of static beams for generating the light sheet. These beams will undergo dithering up and down, meaning that the light sheet will be formed by averaging the profile in the direction of the scan. Notably, the DD optimization disrupts the symmetry of the beam and elongates it in the direction of the scan, a phenomenon that does not impact the Full Width at Half Maximum (FWHM). This optimization demonstrates a sophisticated approach and an physical explanation to the obtained butterfly beam profile.

### 3.3 LSFM imaging results - the butterfly beam improves axial resolution at the edges of the FOV in comparison with Gaussian beam

Next we imaged samples using the butterfly beam, the setup was described in section 2.3. Again, we resized the butterfly phase mask to match the dimensions of the SLM and we multiplied the phase pattern by a constant. The resizing was carried out to optimize the achievable numerical aperture for the system, while the multiplication factor accommodated variations arising from the increasing complexity of the imaging system and the scaling of the phase mask. We have empirically found that a multiplication factor of 160 demonstrated the most significant improvement in the beam profile (Fig. S6).

Figure 4a illustrates the beam profiles in the direction of propagation for both the Gaussian and butterfly beams across the FOV. Additionally, Figure 4a displays the beam waist for the butterfly beam compared to the Gaussian beam at different positions across the FOV. It also provides a comparison with a simulated Gaussian beam, assuming its beam waist matches that of the butterfly beam at the focal plane. This comparison underscores that even when the butterfly beam possesses a larger beam waist, it outperforms a Gaussian beam of the same beam waist.

**Figure 4.**
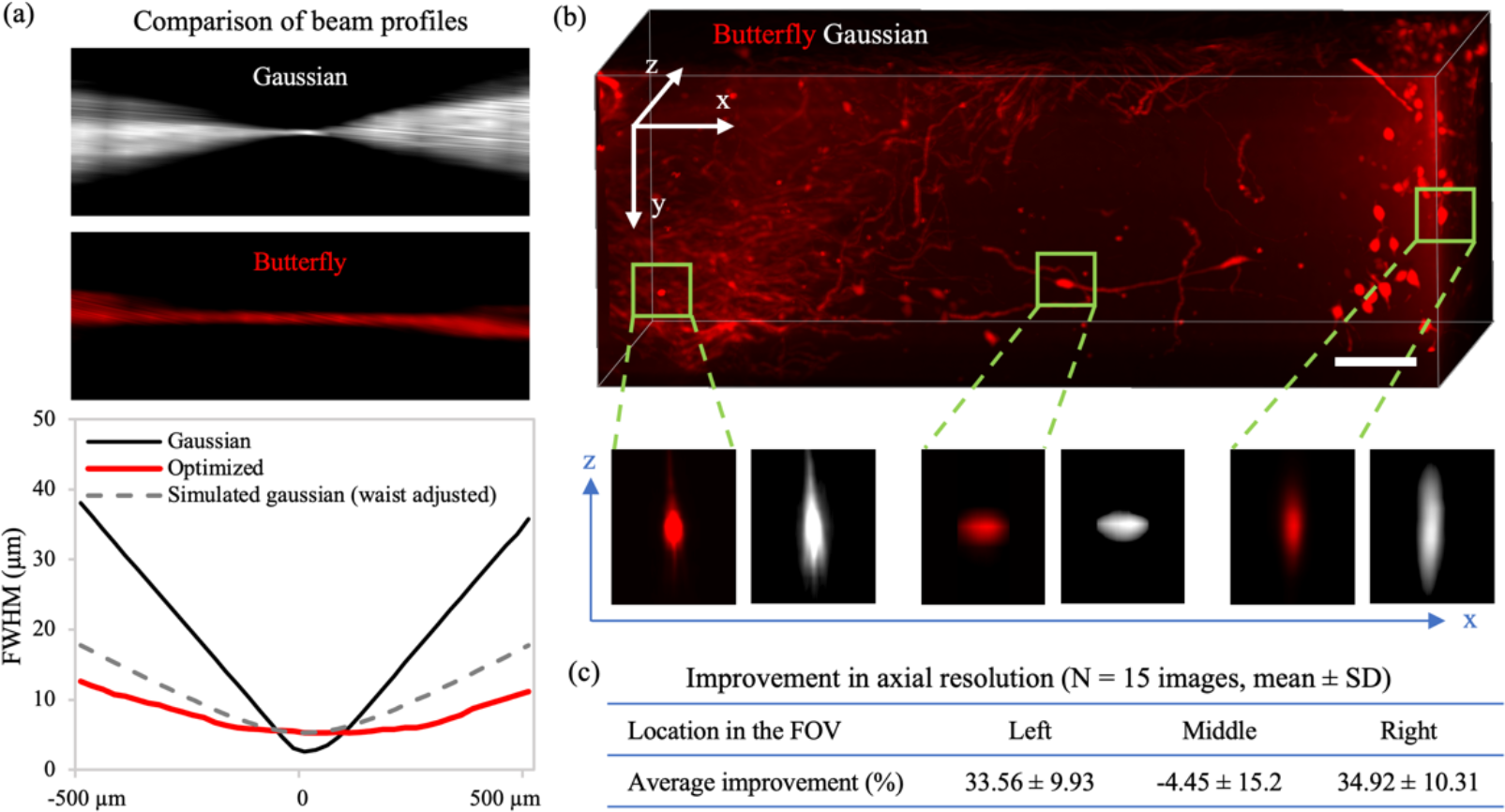
Experimental results for LSFM based imaging. **(a). Comparison of beam profiles**. The beam profile and the FWHM curve demonstrate that the deep design “butterfly” phase mask provides narrower illumination width at the edges of the FOV. Note, the butterfly beam has narrower profile at the edges, even when simulating a Gaussian beam with similar waist. **(b) Maximum intensity projection image of a z-stack acquired from a tissue cleared mouse brain**. The zoomed in images clearly demonstrate that the butterfly beam (red) exhibits a superior axial point spread function (PSF) compared to the Gaussian beam, particularly at the edges. The scale bar is 100 μm. (**c) Improvement in axial resolution**. The butterfly beam has a narrower axial profile in the edges of the FOV whereas in middle the axial profile is comparable to that of the gaussian beam.

In the next series of experiments, we imaged tissue-cleared mouse brain samples in which neurons are sparsely labeled with genetic fluorescent reporters. Figure 4b showcases a 3D volume of a brain sample, with the image obtained using the butterfly beam (red) and the Gaussian beam (white). The axial profiles of several neurons revealed that in the middle region of the FOV (i.e., where the beam waists are similar), both the Gaussian and butterfly beams perform similarly (Fig. 4b, lower half and Fig. S7). However, as we move to the edges of the FOV (i.e., where the butterfly beam exhibits a narrower profile) it became evident that the axial resolution of the butterfly beam surpasses that of the Gaussian beam. We conducted a comparison of the full half maxima in the axial axis for both beams across 15 randomly selected neurons (Fig. 4c) obtained from two brains. The butterfly beam delivered superior optical sectioning at the edges both qualitatively and quantitatively (33-35% improvement in both edges), while the Gaussian beam either matches or provides slightly (∼4.5%) superior results to the butterfly beam in middle, in line with our earlier simulation results.

## 4. Discussion and conclusions

When high-throughput imaging is essential, LSFM emerges as the preferred method, generating extensive imaging datasets commonly analyzed using deep learning models. With the data analysis stage increasingly becoming a bottleneck in utilizing LSFM data, there is a growing demand for approaches that strategically optimize the data acquisition stage to specifically enhance downstream analysis. Several pioneering papers have shown the success of this transformative approach, for instance: deep-STORM was used to enhance super resolution single molecule microscopy, and ‘learned sensing’ was used to improve the detection of malaria infected cells [20,22]. However, in these papers relatively simple optical models were used.

Here, using a complex optical model our novel deep design approach optimized an object detection network and a phase mask that jointly determine the illumination beam shape. This approach is supported by the SLM capability to dynamically modulate the illumination beam, and the ability to simulate the SLM as a large filter in the network structure. Using simulation, DD devised the butterfly beam, and subsequently, we conducted experimental validation, revealing that the butterfly beam exhibits significantly narrower edges within the FOV compared to a Gaussian beam with an identical SLM aperture size. With the narrower beam at the FOV periphery and our objective’s ample depth of field (0.28 NA), we anticipated an immediate enhancement in axial resolution when employing the butterfly beam, as opposed to the Gaussian beam. This observation was consistently validated across tissue cleared brain samples, with the butterfly beam yielding a 33-35% improvement in axial resolution compared to the Gaussian beam.

Upon retrospective examination of the butterfly beam’s shape, a logical explanation emerges. The generation of the light sheet involves dithering the static profile of the beam up and down (Fig. 3b). DD approach capitalizes on this phenomenon, resulting in an asymmetrically elongated butterfly beam along the direction of the scan. Meanwhile, it ensures that the FWHM of the perpendicular axis remains narrow (Fig. 3b). Notably, any elongation or aberrations in the static beam’s direction of the scan fail to impact image quality, as the beam is scanned in this direction. The perpendicular axis (thickness of the light sheet) governs the optical sectioning in LSFM, which the butterfly beam safeguards by maintaining its narrow profile. Upon examination, this approach proves to be physically sound, achieved through pure optimization.

Although other illumination beams such as Bessel and Airy beams can generate similar results, our DD approach can be tailored to any application and optimization goals, if they can be mathematically defined. For instance, DD can be fine-tuned for variable emitter concentrations within a volume or optimized for different loss functions, including increased contrast or resolution, among others. Furthermore, the DD approach presents clear advantages over image enhancement algorithms i.e., post-acquisition enhancement. Unlike such algorithms, which generate intermediary results requiring separate deep learning analysis for information extraction, DD is crafted for end-to-end optimization. The inclusion of image enhancement algorithms adds an extra layer to the analysis process, with no assurance of improved downstream results, as information theory suggests that post-processing alone fails to augment information. In contrast, the DD approach is specifically designed for optimizing the acquisition process to enhance information extraction directly during the image acquisition phase.

An obvious limitation of the presented paper is that we did not use the companion deep learning model to further enhance the image quality. The companion deep learning model (blue pyramid in Fig. 1a) is utilized on down sampled images, resulting in severely degraded resolutions. The reason that we did not employ our companion network on the high-resolution images is that we could not fit the entire image formation model and a large object detection network within our limited 32GB GPU memory. It is crucial to emphasize the indispensable role of GPUs in DD, as their absence results in a tenfold increase in processing time, leading to impractical runtimes per experiment. Given the constant increase in the availability and price of GPUs with higher memory, future employment of a complementary network with the capacity to enhance the performance of the butterfly beam and evaluate its impact on optical sectioning is an immediate future direction for improving our DD approach. It is important to note that the few other groups that have worked on DD, have used a relative rudimentary optical model, and therefore, they could squeeze the optical model into a GPU memory[20,48,49]. An additional limitation of our DD method is that the illumination pattern and the network could converge into local minima, and therefore an optimal solution may be unachievable. We partially addressed this issue by extending our simulation datasets, and by utilizing stochastic gradient descent methods that help to escape local minima.

Overall, the present results constitute a significant stride in exploring novel avenues for system optimization using DD, highlighting the immense potential of our approach to elevate performance across not only imaging pipelines, but other diverse applications.

## Acknowledgments

We would like to thank Andrew Newell and Heather Patisaul for the vole brains samples.

## Data availability

The source code of training end-to-end deep learning model in this paper is not publicly available at this time but may be obtained from the authors upon reasonable request.

