## Supplementary information for "Enhancing Light-Sheet Fluorescence Microscopy Illumination Beams through Deep Design Optimization"

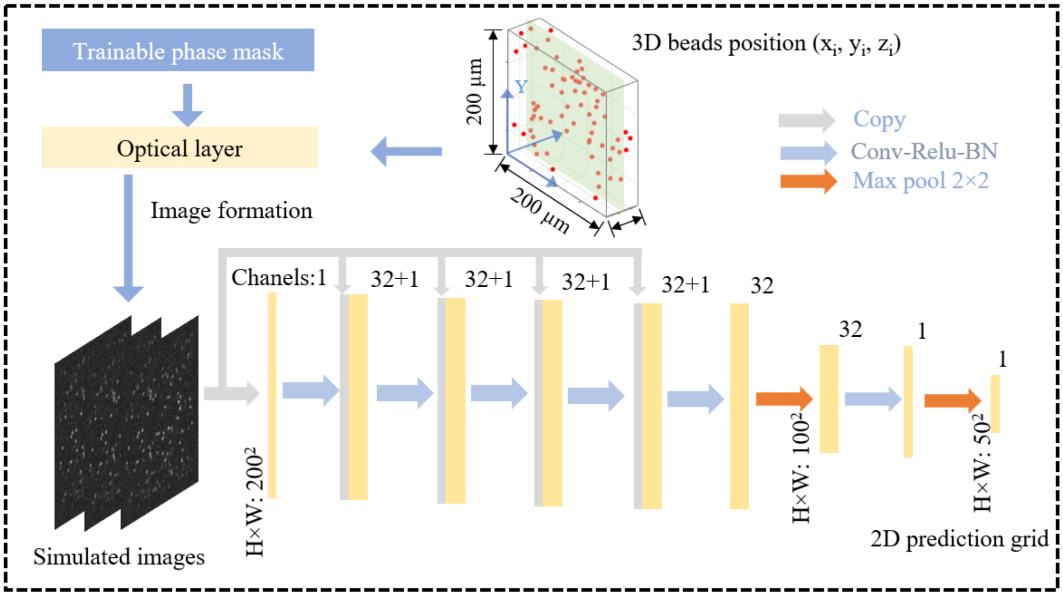


**Fig. S1. Detailed Network structure used for the training.** H – number of pixels in the horizontal direction, W – number of pixels in the vertical direction. Annotation 32+1 - employing 32 filters and a short cut connection.

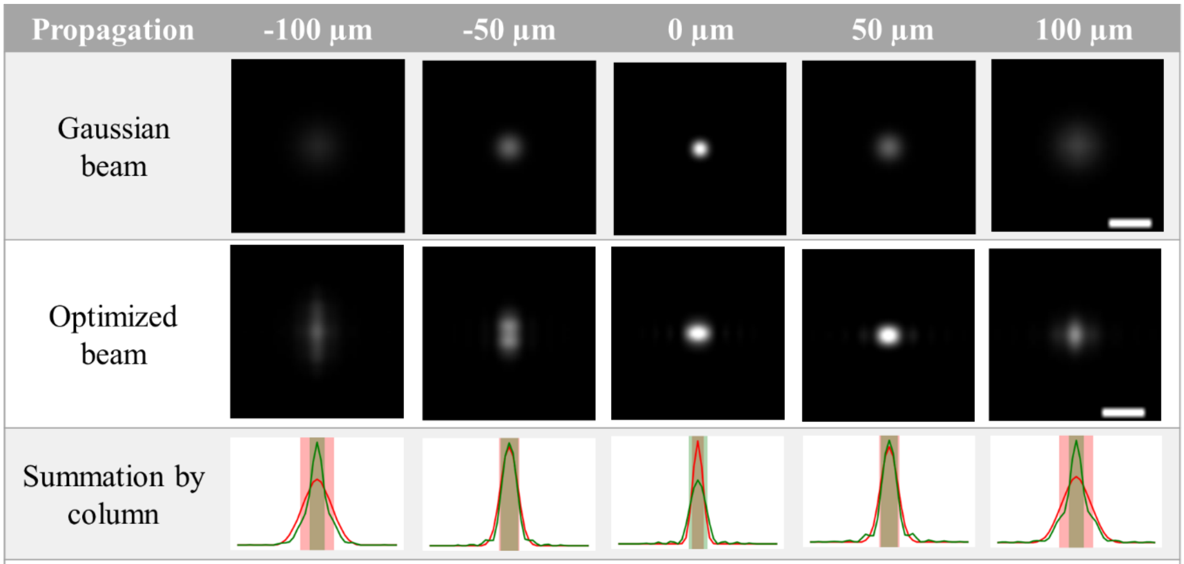


| FWHM of light-sheet (µm) | | | | | |
|---|---|---|---|---|---|
| Gaussian | 8.55 | 5.10 | 3.08 | 5.18 | 8.69 |
| Optimized | 3.87 | 4.43 | 4.72 | 4.32 | 3.87 |

**Fig. S2. Beam profile analysis in simulation**. In the top panels, images depict the static profile of Gaussian and optimized/butterfly beams along the beam propagation. The propagation distance of 0 µm corresponds to the focal point of the excitation objective, where the Gaussian beam reaches its minimal waist. As the light sheet is generated by dithering the static beam up and down, light sheet thickness was measured by summing the values along the column in simulations. Simulation results showcasing that the optimized beam is narrower at the edges of the field of view. The scale bar is 10 µm.

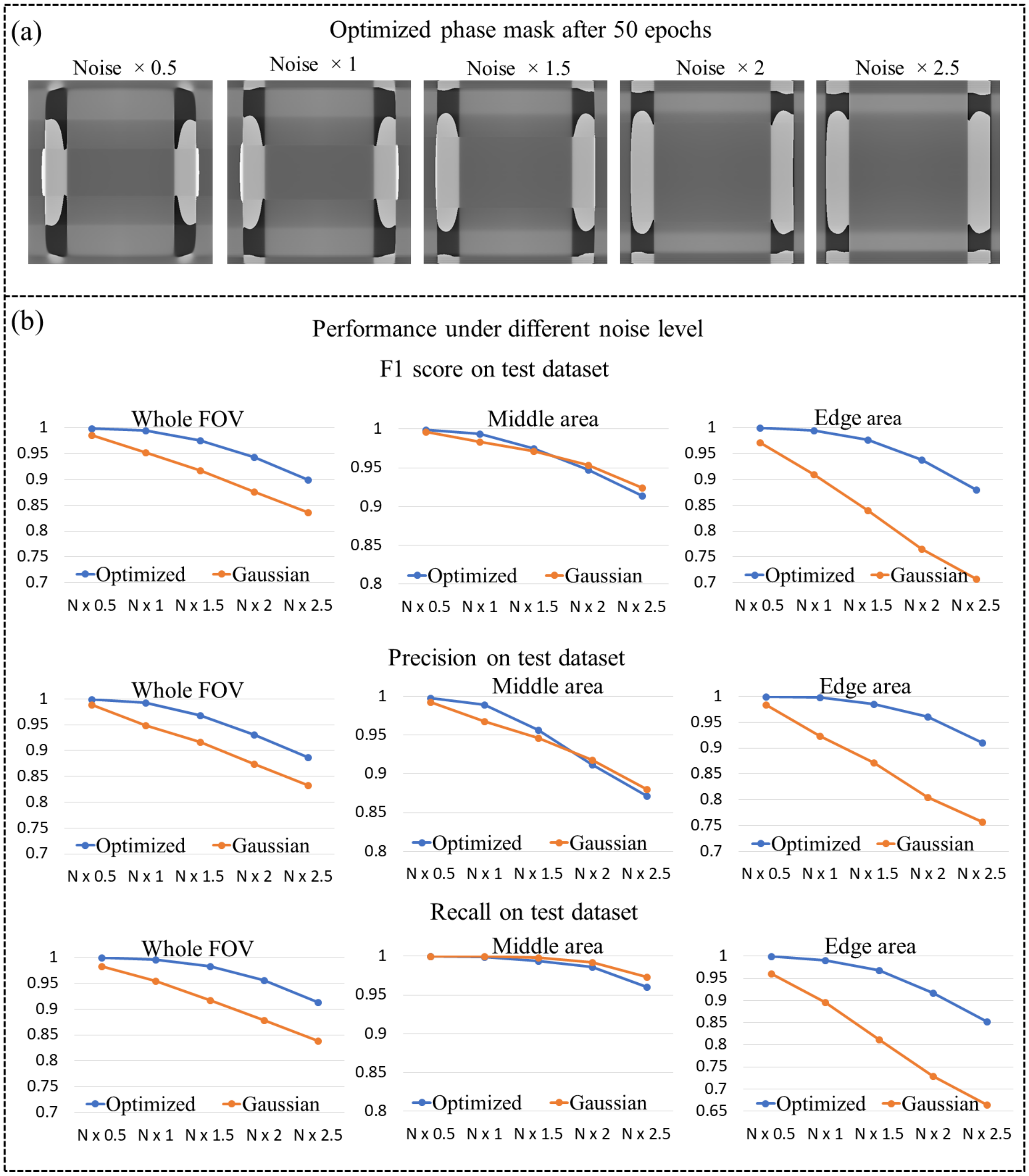


**Fig. S3. Deep Design convergences under varying noise levels in simulation.** (a) The optimized phase after 50 epochs shows consistent patterns despite different levels of Poisson noise, where the standard deviation is multiplied by an increasing factor. This suggests that the butterfly phase pattern generalizes well across varying noise levels. (b) The F1 scores, precision, and recall are depicted under different noise levels using Gaussian beam and butterfly beam on 100 test images across the entire, middle, and edge areas of the field of view. The x-axis indicates the increasing noise level, e.g., $N \times 2$ represents a simulation with double the noise level of $N \times 1$. As anticipated, the butterfly beam performs better than Gaussian under with higher noise levels.

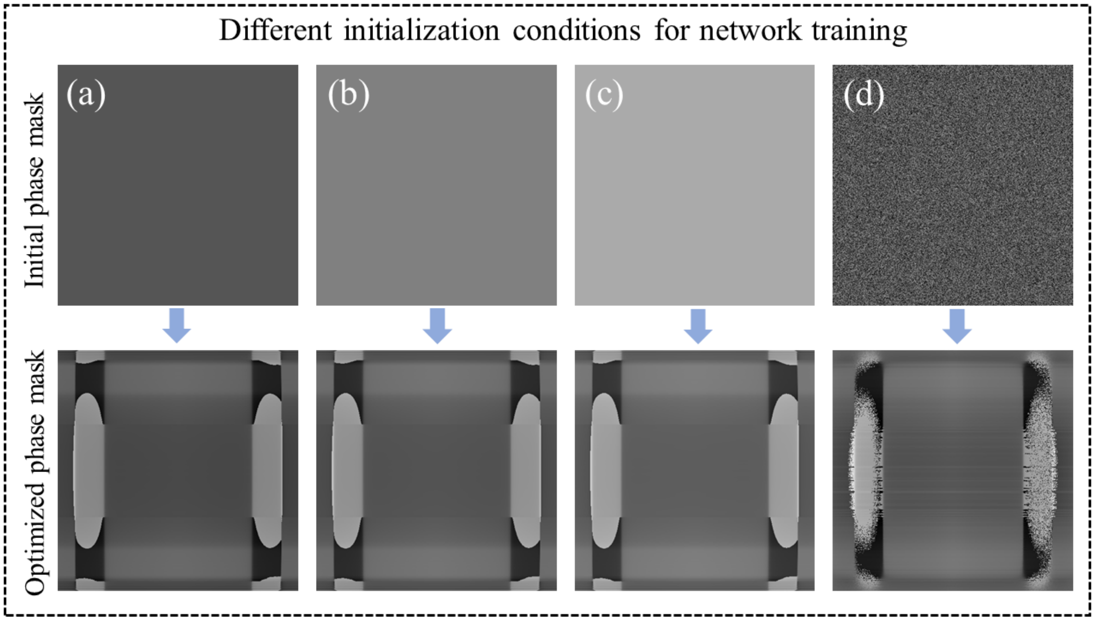


**Fig. S4. Convergence of Deep Design under various initial guesses for the phase mask.** Panels (a)-(d) illustrate initial guesses for the phase mask, beginning with values of all 0, 0.5, 1, and random noise. Despite different initial conditions, all four instances converge to conceptually similar phase masks—ultimately resembling the butterfly phase mask.

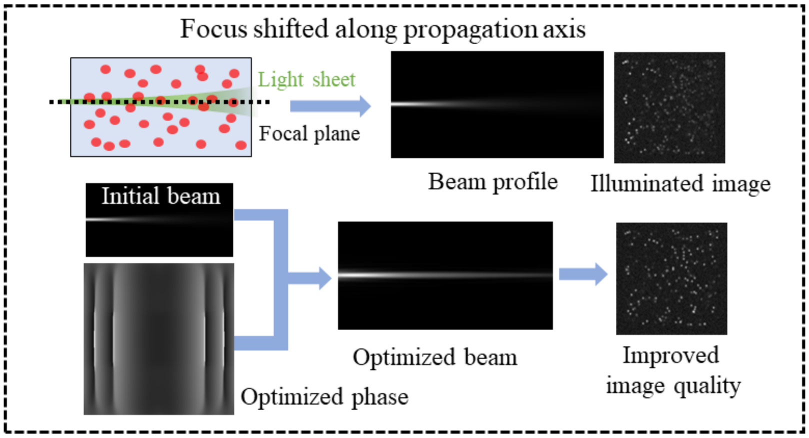


**Fig. S5. Perturbation test for DD.** In instances where the simulation consistently introduces errors in the beam profile—such as focusing on the edge of the field of view under a flat phase mask—DD centers the beam. Conceptually, the resulting phase mask incorporates lens-like patterns atop the butterfly beam. These results underscore the robustness of our approach and its capability to manipulate the light sheet in three-dimensional space.

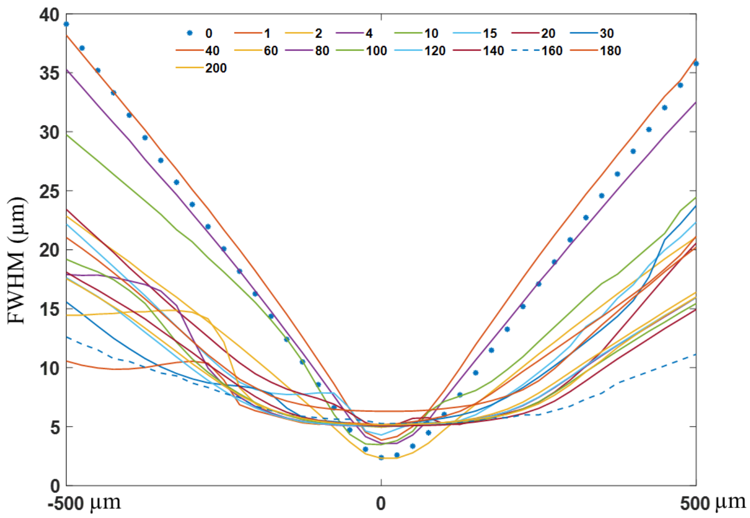


**Fig. S6. In the presence of additional optical elements for imaging tissue cleared samples, the phase mask is multiplied by a modulation factor.** These elements encompass two objective lenses and a sample cuvette filled with immersion media, and they were omitted from the simulation. The Full Width at Half Maximum (FWHM) is assessed for various modulation factors applied to the optimized phase mask. These additional optical elements were not included in the physical model for simplicity but required for imaging tissue cleared samples.

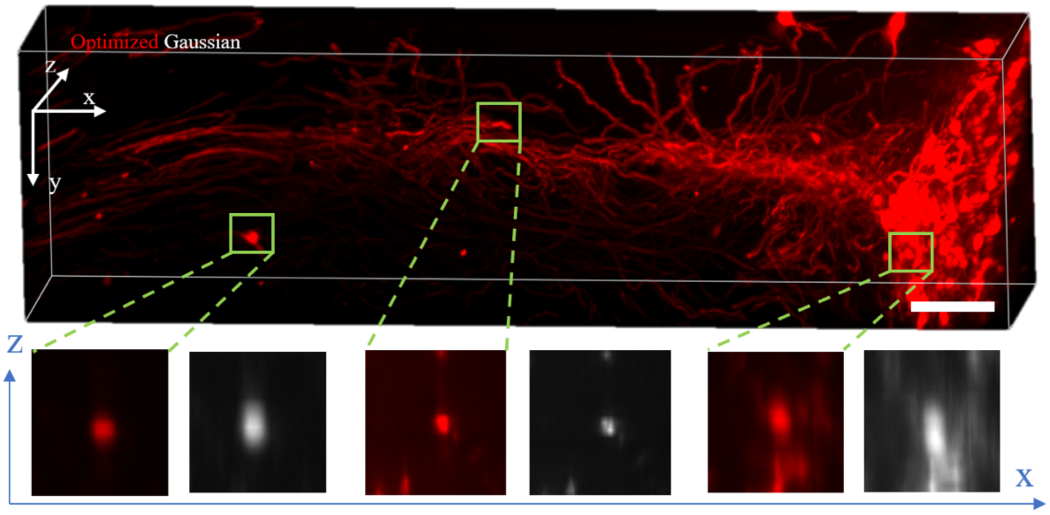


**Fig. S7.** Additional experimental results for LSFM based imaging using the butterfly beam. Maximum intensity projection image of a z-stack acquired from a tissue cleared mouse brain. The zoomed in images clearly demonstrate that the butterfly beam (red) exhibits a superior axial point spread function (PSF) compared to the Gaussian beam, particularly at the edges of the field-of-view. The scale bar is 100 µm.
